# Global Sensitivity Analysis of the Onset of Nasal Passage Infection by SARS-CoV-2 With Respect to Heterogeneity in Host Physiology and Host Cell-Virus Kinetic Interactions

**DOI:** 10.1101/2023.11.04.565660

**Authors:** Leyi Zhang, Han Cao, Karen Medlin, Jason Pearson, Andreas Aristotelous, Alexander Chen, Timothy Wessler, M. Gregory Forest

## Abstract

Throughout the COVID-19 pandemic, positive nasal swab tests have revealed dramatic population heterogeneity in viral titers spanning 6 orders-of-magnitude. Our goal here is to probe potential drivers of infection outcome sensitivity arising from (i) physiological heterogeneity between hosts and (ii) host-variant heterogeneity in the detailed kinetics of cell infection and viral replication. Toward this goal, we apply global sensitivity methods (Partial Rank Correlation Coefficient analysis and Latin Hypercube Sampling) to a physiologically faithful, stochastic, spatial model of inhaled SARS-CoV-2 exposure and infection in the human respiratory tract. We focus on the nasal passage as the primary origin of respiratory infection and site of clinical testing, and we simulate the spatial and dynamic progression of shed viral load and infected cells in the immediate 48 hours post infection. We impose immune evasion, i.e., suppressed immune protection, based on the preponderance of clinical evidence that nasal infections occur rapidly post exposure, largely independent of immune status. Global sensitivity methods provide the de-correlated outcome sensitivities to each source of within-host heterogeneity, including the dynamic progression of sensitivities at 12, 24, 36, and 48 hours post infection. The results reveal a dynamic rank-ordering of the drivers of outcome sensitivity in early infection, providing insights into the dramatic population-scale outcome diversity during the COVID-19 pandemic. While we focus on SARS-CoV-2, the model and methods are applicable to any inhaled virus in the immediate 48 hours post infection.

## 1 Introduction

The COVID-19 pandemic has amplified the critical need to understand the mechanistic drivers of diverse outcomes in the dynamic and spatial progression of viral infection *within* the human respiratory tract (RT), and thereby shed insights on the observed dramatic population heterogeneity in infection outcomes. In particular, clinical nasal swab titers of non-hospitalized infected individuals have varied over the pandemic by 6 orders of magnitude [1, 2, 3, 4, 5, 6, 7, 8, 9]. Since January of 2020, our group has developed multi-scale (in space and time), within-host, human RT infection models and the necessary software tools to simulate this intricate process. The baseline model [10] incorporates the complex anatomy and physiology of the human RT, as well as the kinetic processes of virion diffusivity *D*_*v*_ in the mucosal barrier, the probability *p*_infect_ of cell infection per virion encounter-second, the latency period *t*_latency_ of an infected cell, the shedding rate *r*_shedding_ of infectious viral RNA copies, and duration of the shedding phase until cell death. The latency time *t*_latency_ spans a positive infectious encounter until onset of extracellular shedding of viral RNA, which covers cellular uptake of the virus and viral hijacking of cellular replication to produce RNA copies. Typically the time from initial infection of a cell to its death is longer than 48 hours, so this factor will not be relevant to this study. The parameters of the model implemented in [10] used mean population properties of human RT physiology (e.g., for each branch generation in the upper and lower RT, branch length, mucus thickness, and mucociliary advection velocity of the mucus layer) together with the best-known clinical and experimental parameter estimates of the kinetic processes for SARS-CoV-2 virions. The alveolar ducts and sacs of the deep lung were modeled as well in [11], but those details are not relevant for the present focus on nasal infections.

In [12], the sensitivity of viral load and infected cells in the nasal passage due to variations in several infection-related parameters was explored. Sensitivity due to physiological heterogeneity was not explored; all model simulations were performed for fixed mucus thickness and advection velocity at mean population estimates. While insightful with respect to outcome sensitivity, the analysis in [12] was based on fixing all other parameters at mean estimates while varying each individual parameter over a potential range of values. Here we replace the analysis based on these “1-dimensional slices of parameter space” with a random sampling of the full parameter space to gain a more faithful measure of sensitivities. Several insights from [12] guide the present study with regards to the early outcomes of shed viral load and number of infected cells. First, it was discovered that early outcomes are extremely robust to variations in *p*_infect_, and as a consequence we choose to fix *p*_infect_ = 0.2. Second, early outcomes were found to be sensitive to *r*_shedding_. However, the experimental and clinical data on the replication rate of *infectious* RNA copies remains poorly understood, and so we explore two decades of *r*_shedding_, 10 − 1000 virions/day. Finally, early outcomes were seen to be exponentially sensitive to linear scaling of *t*_latency_, and so based on prior [13, 14] and continued [15] single-cell experimental resolution data we explore *t*_latency_ spanning 3 − 9 hours. (N.B. Since we fix *p*_infect_ = 0.2 in this analysis, results from [12] are presented in Supplementary Material to illustrate the remarkable robustness of outcomes to an order of magnitude variability in *p*_infect_.)

Upper and lower bounds on all parameters, both physiological and in virus-cell infection kinetics, continue to be updated during the pandemic. Remarkably, none of the three cellular kinetic parameters in our model have been explicitly quantified, so we retain bounds on the sensitive parameters *r*_shedding_ and *t*_latency_ that are consistent with the literature noted just above, and fix the robust kinetic parameter *p*_infect_ = 0.2. There is, however, strong clinical and experimental evidence [16] that two physiological parameters vary significantly with SARS-CoV-2 infection: the thickness *M*_thickness_ and the mucociliary advection velocity *M*_vel_ of the mucus layer in the nasal passage. To our knowledge, the impact of host heterogeneity in these fundamental physiological features of nasal infection has never been explored, not just for SARS-CoV-2, but for any virus. Therefore, in this study we explore sensitivity of the dynamic outcomes over 48 hours in infectious viral load, total number of infected cells, and flux of infectious viral RNA copies out of the nasal passage, globally across the four-parameter space [*r*_shedding_, *t*_latency_, *M*_thickness_, *M*_vel_].

Our baseline model assumes the absence of all forms of immune protection over the first 48 hours. This assumption was obviously valid at the onset of the pandemic throughout the human population, and continues to apply to the immuno-compromised, unvaccinated, and previously uninfected. Yet remarkably, the continuing clinical cases of infection by vaccinated and previously infected persons throughout the waves of dominant variants [17, 18, 19, 20, 21, 22, 23,24, 25, 9] reveal that immune escape and evasion by SARS-CoV-2 persist for 48 hours or longer, independent of vaccination status and prior infection. For this reason, there is much to be learned about the driving factors of nasal passage infection in the immediate 48 hours post infection from the pre-immune RT infection model.

## 2 The Model

As shown in Figure 1 from [10], and articulated in detail in [26], the nasal passage and all generations of the lower RT except the alveolar space are approximately cylindrical. In each generation, the epithelial cell surface is coated by a 7 *μ*m thick layer of periciliary liquid (PCL) in which cilia beat. At full extension in the power stroke, cilia penetrate the PCL-mucus interface and extend into the mucus layer about 1 *μ*m, and the coordinated metachronal waves of cilia propel the mucus layer, “down” in the nasal passage and “up” in the lower RT, towards the esophagus to be swallowed.

**Figure 1.**
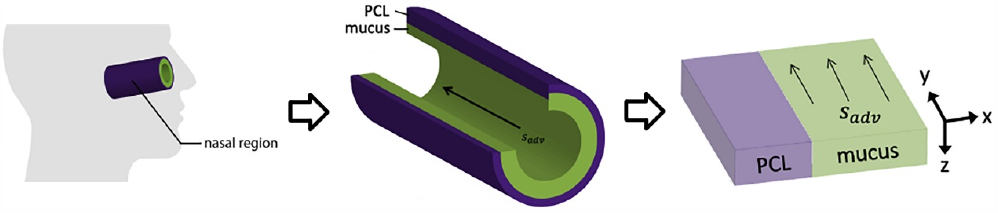
Modeling the nasal passage (image taken from [10]).

We unfold this cylindrical geometry into a rectangular domain in which the *y*-*z*-plane falls on the epithelial cell surface. *x* denotes the “radial” distance into the PCL and mucus layers, with *x* = 0 being the epithelium-PCL interface. *y* denotes the distance along the centerline axis, which is the primary direction of mucus advection by coordinated beating of cilia, with *y* = 0 representing the entry into the nasal passage. *z* is the azimuthal axis of the cylinder. Infectious virions undergo diffusion in PCL and mucus, and additional advection with velocity *M*_vel_ while in the mucus layer, governed by:

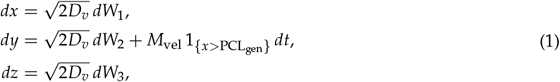

Where

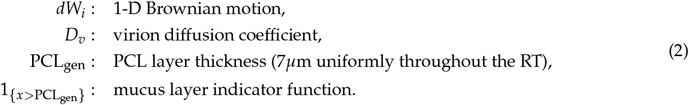

Ciliated cells are the predominant infectable cells in the RT above the alveolar space, covering about 50% of the epithelial surface. Every epithelial cell has a degree of infectability, either non-infectable or with a prescribed probability *p*_infect_ of infection per encounter-second.

In our model, a freely diffusing virion in the PCL encounters a cell when its distance from the epithelial cell surface vanishes, i.e. when *x* = 0. For each second during which an encounter with a ciliated cell takes place, there is a probability *p*_infect_ of an infection. If an encounter results in infection, the cell switches from uninfected to infected, and the virion is removed from the free virion population. When the stochastic virus-cell encounter does not result in infection, for infectable or non-infectable cells, the virion is reflected back into the PCL.

Each virion is tracked until it either infects a cell or exits the generation, always toward the trachea due to strong mucus advection. Once a cell switches to an infected state, it persists in an infected, non-shedding latency state for a duration *t*_latency_, which represents cellular uptake of the virus and hijacking of the cellular machinery to replicate viral RNA copies. After *t*_latency_ has lapsed, the cell switches to a shedding state, replicating infectious virions at rate *r*_shedding_. Since infected cells typically die after 3 days post infection, no cells switch to a death state in this 48-hour study.

We assume that the kinetics of SARS-CoV-2 interactions with ciliated cells are robust within each host, yet potentially highly variable between hosts, and therefore we explore literature-supported ranges for the kinetic parameters that our previous study [12] revealed to be sensitive. All simulations to generate data for this study start at the moment of infection of one cell at the entry of the nasal passage (axial coordinate *y* = 0). Table 1 summarizes the model parameters, fixed and variable, and the simulation details. Table 2 summarizes the three model outcomes and associated data.

**Table 1:**
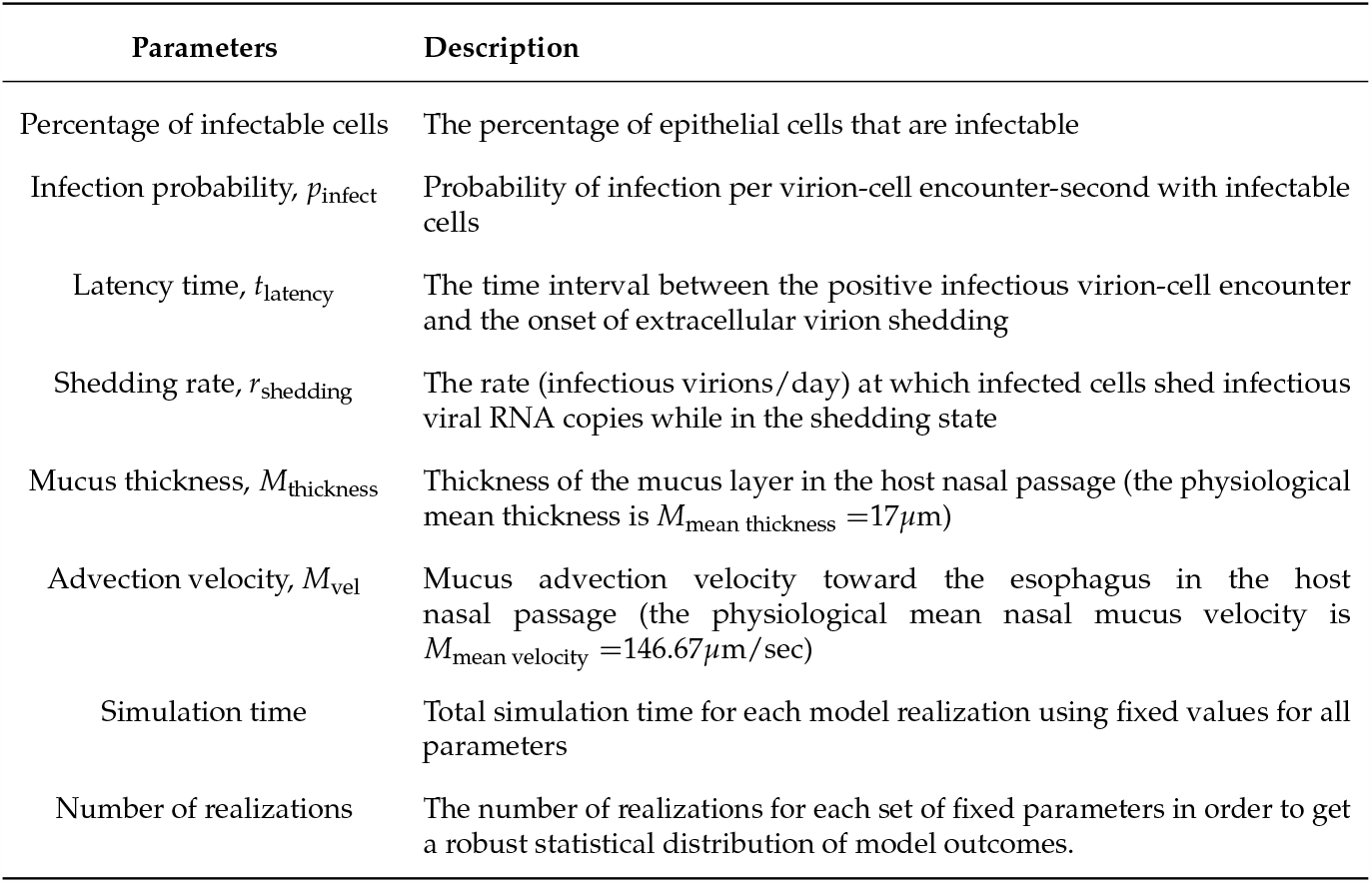
The model parameters and their descriptions.

**Table 2:** The model outcomes and their descriptions.

| Outcome Data | Description |
| --- | --- |
| Total viral load | The number of freely diffusing infectious virions in the nasal passage |
| Infected cell count | The total number of cells that have been infected |
| Virion Flux | The number of infectious virions that have exited the nasal passage via mucus transport |

## 3 Methods

In [12], we explored the sensitivity of outcomes from an initial nasal infection to host cell-virus kinetic parameters. In that study, one parameter was varied across an estimated range of possible values, while all other parameters were fixed at best-known mean estimates. Some key results are: the total numbers of infected cells and total viral load are remarkably robust to variations in cell infectivity, *p*_infect_; shorter latency time *t*_latency_ has a dramatic, exponential effect on the progression of infected cells and total viral load; and, higher shedding rate *r*_shedding_ of infected, post-latency cells has a significant proportional (yet non-exponential) effect on infected cell count and total viral load.

While insightful, these sensitivities correspond to 1-dimensional slices in the parameter space being explored, and therefore lack the ability to detect if the sensitivities gained are robust to sampling off that 1-dimensional slice. To generalize these limited searches of parameter space requires methods that perform global sampling and sensitivity analysis, which we summarize next and then apply. Further, our previous studies did not explore host-to-host physiological heterogeneity, which recent studies [16] have shown to arise during SARS-CoV-2 infection. We therefore add two physiological parameters, mucus thickness and advection, to our global sampling and sensitivity analyses.

### 3.1 Latin Hypercube Sampling

Latin Hypercube Sampling (LHS) is a widely used technique to sample high-dimensional parameter spaces. It offers a quasi-random approach to efficiently sample across the entire parameter space while minimizing the number of required sampling points. Implementing LHS allows exploration of a wide range of parameters at a high resolution.

In addition, we apply Partial Rank Correlation Coefficient (PRCC) to the simulated data, which we will introduce in Section 3.2. The sampling strategy of [12] contains repeated parameter values, which can impact the accuracy of PRCC results, so we cannot directly apply PRCC. Implementing LHS alleviates this issue.

LHS can be carried out as follows:

1. Start by selecting the sample size *N*. This will be the number of our sample points in the parameter space.
2. Determine the range and distribution of each parameter (e.g. we chose a uniform distribution for *t*_latency_ ranging from 3 hours to 9 hours).
3. Divide the range of each parameter into *N* equal-probability intervals
4. Repeat the following steps *N* times:
  a. For each parameter, randomly select one interval from the remaining pool of intervals
  b. Randomly sample from the selected intervals for all parameters
  c. Remove the selected intervals from the remaining pool of intervals

Figure 2 shows an example of using LHS with sample size *N* = 20 on two parameters (latency time and advection velocity with uniform distributions). We see that the range of each parameter is evenly divided into 20 intervals. Each column and each row contains exactly one sample point.

**Figure 2.**
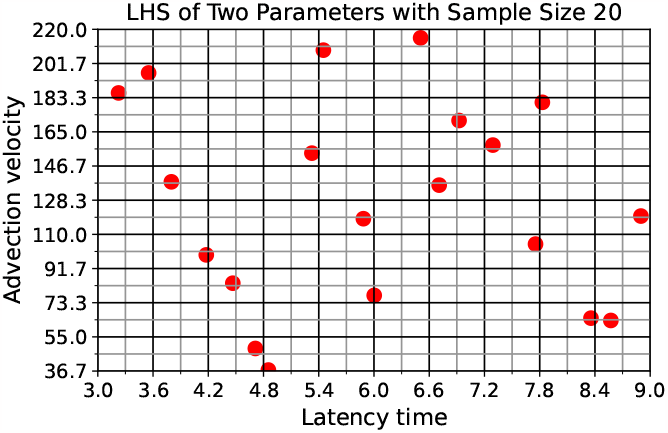
An example in which LHS is applied on two parameters using sample size 20.

Table 3: shows the parameter ranges and distributions chosen for this analysis.

**Table 3:**
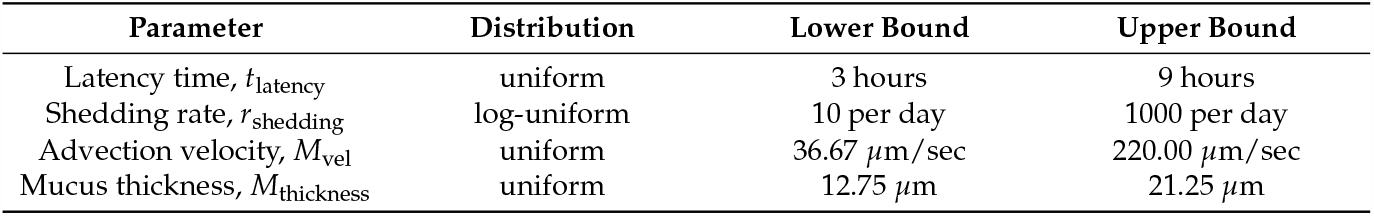
The 4-dimensional parameter space for the PRCC sensitivity analysis. For mucus advection velocity and mucus thickness, we apply a range of multiplicative factors to the population mean average values from [10]. For mucus advection velocity *M*_vel_, we choose the range to be .75 to 1.25 times the physiological mean thickness at 17*μ*m. For mucus thickness, we choose the range to be .25 to 1.25 times the physiological mean nasal mucus velocity at 146.67*μ*m/sec.

### 3.2 Partial Rank Correlation Coefficient Method

The Partial Rank Correlation Coefficient Method, or PRCC, is a sensitivity analysis technique that provides a measure of the correlation between individual parameters and model outcomes, whereby effects from cross-correlations with all other variables are calculated and removed [27]. The implementation of PRCC starts with rank transforming the correlation parameters *x*_*j*_ and outcomes *y*. For each index *j*, we perform linear regression on *x*_*j*_ and *y* in terms of other parameters:

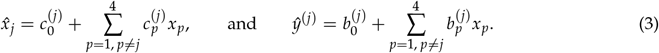

The PRCC is the Pearson Correlation Coefficient (PCC) between the residuals, 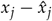 and *y* − *ŷ*^(*j*)^, given by

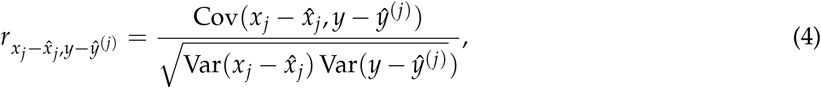

where 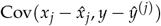 represents the covariance between the residuals, and 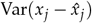and Var(*y* − *ŷ*^(*j*)^) represent the variance of 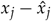 and *y* − *ŷ*^(*j*)^, respectively.

The resulting PRCC value for each parameter is a number between − 1 and 1, where the sign indicates positive or negative correlation and the magnitude indicates the degree of sensitivity of the outcome in question to variations in the parameter.

### 3.3 Simulations and Data Generation

Prior to the sensitivity analysis step, we sampled the four-parameter space using LHS as described in section 3.1 with sample size *N* = 20. As shown in Table 4, we fix the value *p*_infect_ = 0.2 based on the results in [12] showing extreme robustness in outcomes over a decade or more variations. We also fix the percentage of infectable cells at 50% corresponding to the percentage of ciliated cells; this value could be slightly higher, but again the outcomes are robust to variations [12]. We record the total shed infectious viral load, the infected cell count, and the viral flux from the nasal passage at 12, 24, 36, and 48 hours post infection of a single cell at the entry of the nasal passage.

**Table 4:** These parameter values are fixed for this study.

| Parameters | Values |
| --- | --- |
| Infection probability, $p_{\text{infect}}$ | 0.2 |
| Percentage of infectable cells | 50% |
| Simulated time | 48 hours |
| Number of realizations | 75 |

### 3.4 An Example

We use the infected cell count at 36 hours as an example to demonstrate how we compute the PRCC analysis applied to simulation data from our spatial nasal infection model. Recall that we have previously established the following notations to represent the parameters, and we choose *y* to denote the outcomes.

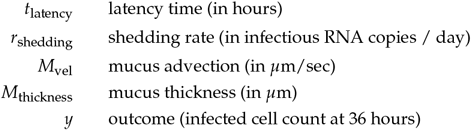

The parameter columns in Table 5a show all 20 points in the four-dimensional parameter space selected by the LHS process. Each row represents a parameter combination and the corresponding mean simulation outcome. Figure 3 shows scatter plots of the outcome values verses each parameter.

**Table 5:** Parameter values and example outcome data: (a) shows all 20 parameter combinations selected by the LHS process, corresponding to the infected cell count at 36 hours; (b) shows the rank transformed parameter values and the outcome data from a.

| Parameters |  |  |  | Outcomes |
| --- | --- | --- | --- | --- |
| $t_{\text{latency}}$ | $r_{\text{shedding}}$ | $M_{\text{vel}}$ | $M_{\text{thickness}}$ | $y$ |
| 6.71 | 17 | 136.41 | 18.66 | 269 |
| 5.88 | 217 | 118.40 | 18.18 | 683,012 |
| 6.51 | 111 | 215.41 | 16.44 | 38,445 |
| 3.55 | 174 | 196.61 | 14.97 | 1,167,785 |
| 5.33 | 833 | 153.57 | 13.21 | 1,546,895 |
| 8.36 | 10 | 65.06 | 14.24 | 62 |
| 6.92 | 41 | 171.09 | 17.77 | 1,759 |
| 8.90 | 25 | 119.83 | 21.22 | 200 |
| 3.23 | 132 | 185.87 | 15.80 | 1,154,757 |
| 4.47 | 78 | 83.81 | 14.63 | 505,501 |
| 7.75 | 14 | 104.79 | 19.00 | 123 |
| 4.71 | 457 | 48.80 | 16.63 | 2,280,089 |
| 6.00 | 88 | 77.26 | 20.00 | 79,233 |
| 4.18 | 353 | 99.00 | 19.16 | 1,760,488 |
| 5.45 | 39 | 208.82 | 15.64 | 4,642 |
| 4.85 | 582 | 37.22 | 20.52 | 2,615,026 |
| 8.57 | 793 | 63.81 | 12.91 | 706,248 |
| 3.80 | 24 | 138.19 | 17.22 | 8,444 |
| 7.29 | 58 | 157.90 | 13.80 | 4,426 |
| 7.83 | 268 | 180.84 | 19.64 | 105,808 |

| Parameters (ranks) |  |  |  | Outcomes (ranks) |
| --- | --- | --- | --- | --- |
| $t_{\text{latency}}$ | $r_{\text{shedding}}$ | $M_{\text{vel}}$ | $M_{\text{thickness}}$ | $y$ |
| 13 | 3 | 11 | 14 | 4 |
| 10 | 14 | 9 | 13 | 13 |
| 12 | 11 | 20 | 9 | 9 |
| 2 | 13 | 18 | 6 | 16 |
| 8 | 20 | 13 | 2 | 17 |
| 18 | 1 | 4 | 4 | 1 |
| 14 | 7 | 15 | 12 | 5 |
| 20 | 5 | 10 | 20 | 3 |
| 1 | 12 | 17 | 8 | 15 |
| 5 | 9 | 6 | 5 | 12 |
| 16 | 2 | 8 | 15 | 2 |
| 6 | 17 | 2 | 10 | 19 |
| 11 | 10 | 5 | 18 | 10 |
| 4 | 16 | 7 | 16 | 18 |
| 9 | 6 | 19 | 7 | 7 |
| 7 | 18 | 1 | 19 | 20 |
| 19 | 19 | 3 | 1 | 14 |
| 3 | 4 | 12 | 11 | 8 |
| 15 | 8 | 14 | 3 | 6 |
| 17 | 15 | 16 | 17 | 11 |

(a) Parameter values and outcome values
(b) The ranks of parameter values and the ranks of outcome values

**Figure 3.**
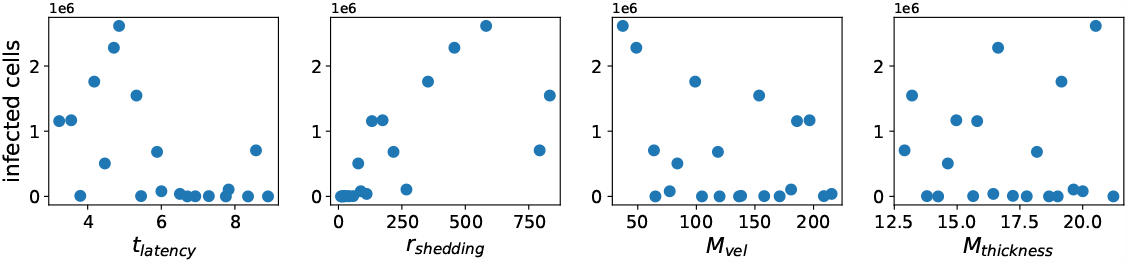
Infected cell count at 36 hours versus each parameter.

Within each column of Table 5a, we rank-transform the column by assigning integers from 1 to 20 to values ranking from the smallest to the largest. Table 5b shows the rank-transformed parameters and outcome values. Figure 4 shows scatter plots of the ranks of outcome values verses the ranks of each parameter.

**Figure 4.**
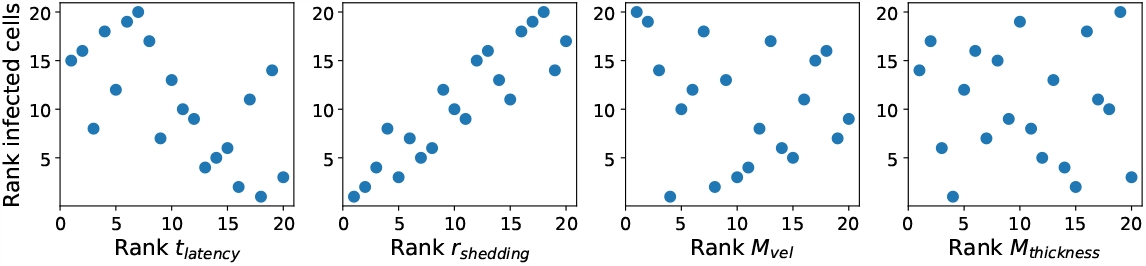
Ranks of infected cell count at 36 hours versus ranks of each parameter. Integers from 1 to 20 are assigned to values ranking from the smallest to the largest.

Then we perform linear regression on each rank-transformed parameter and outcome in terms of the other parameters.

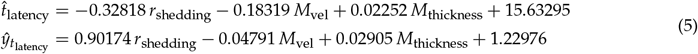

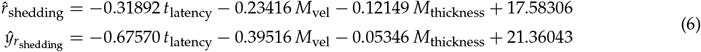

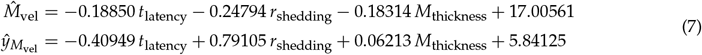

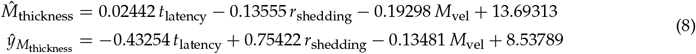

Finally, for each parameter *x* ∈*{t*_latency_, *r*_shedding_, *M*_vel_, *M*_thickness_*}*, we compute the PCC between the residuals 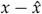 and *y*− *ŷ*_*x*_ using the formula in Eq. 4. The resulting numbers are the PRCCs between the parameters and the outcomes. Figure 5 shows scatter plots of residuals of rank-transformed outcome values versus residuals of rank-transformed parameters.

**Figure 5.**
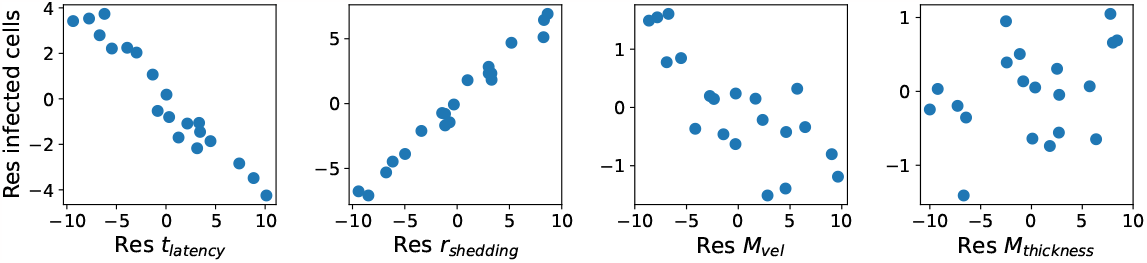
Residuals of infected cell count at 36 hours versus the residuals of each parameter. The residuals are produced by subtracting linear regression models from the outcome ranks and the parameter ranks. The CC between the residuals is the PRCC.

Table 6 shows a comparison between the Pearson Correlation Coefficients (PCC), the Spearman Correlation Coefficients (SCC), and the Partial Rank Correlation Coefficients (PRCC) for each parameter. Note that SCC are obtained by computing the PCC after rank transforming the data (as in Table 5b and Figure 4), while PRCC are obtained by computing the PCC after rank transforming the data and taking the residuals of the data (as in Figure 5).

**Table 6:** Comparisons of PCC, SCC, and PRCC for infected cell count at 36 hours for all parameters.

| Parameters | PCC | SCC | PRCC |
| --- | --- | --- | --- |
| Latency time, $t_{\text{latency}}$ | -0.54573 | -0.64060 | -0.97203 |
| Shedding rate, $r_{\text{shedding}}$ | 0.69960 | 0.90677 | 0.99036 |
| Advection velocity, $M_{\text{vel}}$ | -0.40211 | -0.20752 | -0.77671 |
| Mucus thickness, $M_{\text{thickness}}$ | -0.00101 | -0.6165 | 0.35999 |

Scatter plots similar to Figures 3, 4, 5 showing (1) raw outcome data vs. parameter values, (2) ranks of outcome data vs. ranks of parameter values, and (3) residuals of outcome data vs. residuals of parameter values, for all three outcomes (total viral load, infected cell count, and flux) at 12-hour time increments (12, 24, 36, 48 hours) can be found in Appendix B.

## 4 Results

In the implementation, we use the R function epi.prcc() from the epiR package to compute PRCC between each parameter for each of the the three types of outcome data (shed infectious virion count, infected cell count, infectious virion flux via mucus clearance) at 12, 24, 36, and 48 hours following the initial nasal cell infection at the entry of the nasal passage.

We observe that latency time *t*_latency_ and extracellular shedding rate of virions *r*_shedding_ have a significant impact on all infection outcomes at all timestamps. The influence of mucus advection velocity *M*_vel_ progressively intensifies from weak to somewhat strong for the total shed viral load and infected cell count as time progresses over the first 48 hours post initial cell infection. Mucus thickness *M*_thickness_ within these physiological bounds has a relatively minor impact on all infection outcomes.

Figure 6 and Table 7 show PRCC results for *total viral load* for all four parameters at four 12-hour time increments over 48 hours post infection. With extremely high likelihood, independent of other parameter choices, lower values of *t*_latency_ within the 3-9 hour range exponentially increases total viral load at all timestamps. Similarly increasing *r*_shedding_ over a logarithmic range of 10 to 1000 infectious virions per day induces an exponential increase in total viral load at all timestamps. Slower mucus advection, as reported in [16] for COVID-19 infection, amplifies the total viral load and the number of infected cells (shown in Figure 7 and Table 8), with the effect becoming stronger over the 48 hours post infection. We do not detect a significant effect of mucus thickness.

**Table 7:** PRCC results for total viral load at 12, 24, 36 and 48 hours for model parameters: latency time (*t*_latency_), extracellular shedding rate of infectious RNA copies (*r*_shedding_), mucus advection velocity, and mucus thickness.

|  | 12 Hours | 24 Hours | 36 Hours | 48 Hours |
| --- | --- | --- | --- | --- |
| Latency time, $t_{\text{latency}}$ | -0.94123 | -0.97044 | -0.97263 | -0.96750 |
| Shedding rate, $r_{\text{shedding}}$ | 0.97881 | 0.99009 | 0.99181 | 0.99327 |
| Advection velocity, $M_{\text{vel}}$ | -0.56282 | -0.61626 | -0.71987 | -0.83138 |
| Mucus thickness, $M_{\text{thickness}}$ | 0.19907 | 0.37230 | 0.10946 | -0.17701 |

**Table 8:** PRCC results for infected cell count at 12, 24, 36 and 48 hours for model parameters: latency time (*t*_latency_), extracellular shedding rate of infectious RNA copies (*r*_shedding_), mucus advection velocity, and mucus thickness. Identical PRCC can occur due to identical ranks.

|  | 12 Hours | 24 Hours | 36 Hours | 48 Hours |
| --- | --- | --- | --- | --- |
| Latency time, $t_{\text{latency}}$ | -0.96017 | -0.97203 | -0.97203 | -0.95666 |
| Shedding rate, $r_{\text{shedding}}$ | 0.98461 | 0.99036 | 0.99036 | 0.98748 |
| Advection velocity, $M_{\text{vel}}$ | -0.70904 | -0.77671 | -0.77671 | -0.86441 |
| Mucus thickness, $M_{\text{thickness}}$ | 0.27081 | 0.35999 | 0.35999 | -0.10695 |

**Figure 6.**
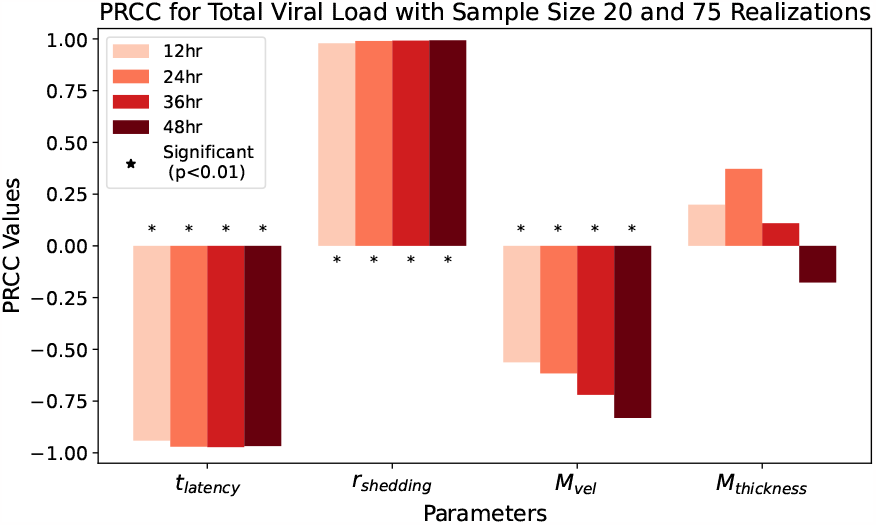
PRCC results for total viral load at 12, 24, 36, and 48 hours.

**Figure 7.**
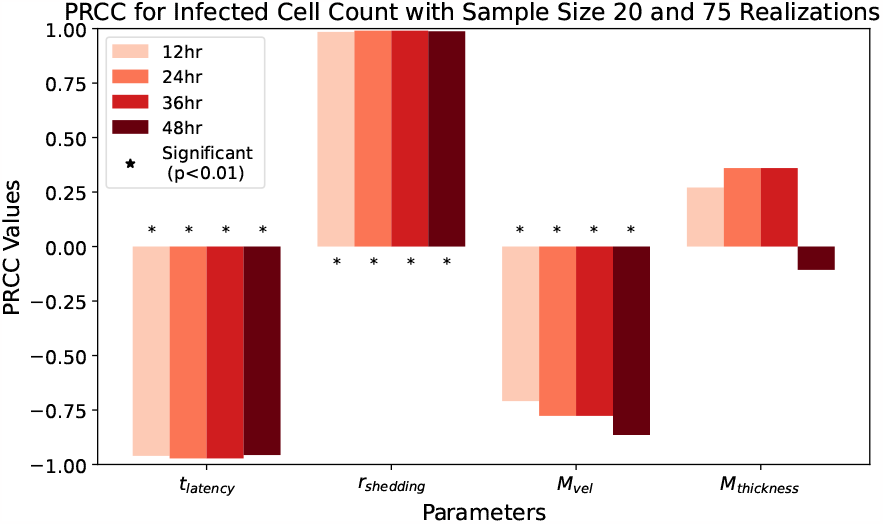
PRCC results for infected cell count at 12, 24, 36, and 48 hours.

Figure 7 and Table 8 show PRCC results for *infected cell count* at 12-hour time increments for the selected parameters. The results look very similar to those in Figure 6 and Table 7, except that the effect of *M*_vel_ has a weaker time dependence, with the effect being more noticeable earlier in the infection compared to its effect on total viral load.

Note that in Table 8, the values in column “24 Hours” match up exactly to those in column “36 Hours”. Investigation of the data shows that the ranks of the infected cell counts were preserved from 24 hours to 36 hours, while the raw data values changed over time. The identical PRCC values are a consequence of the identical ranks.

Figure 8 and Table 9 show PRCC results for *viral flux* (the total number of virions transported out of the nasal passage via mucus advection). Similar to previous results, *t*_latency_ has a strong negative correlation with flux and *r*_shedding_ has a strong positive correlation with flux.

**Table 9:** PRCC results for virion flux at 12, 24, 36 and 48 hours for model parameters: latency time (*t*_latency_), extracellular shedding rate of infectious RNA copies (*r*_shedding_), mucus advection velocity, and mucus thickness.

|  | 12 Hours | 24 Hours | 36 Hours | 48 Hours |
| --- | --- | --- | --- | --- |
| Latency time, $t_{\text{latency}}$ | -0.85576 | -0.91655 | -0.93570 | -0.96692 |
| Shedding rate, $r_{\text{shedding}}$ | 0.83963 | 0.94356 | 0.97463 | 0.98938 |
| Advection velocity, $M_{\text{vel}}$ | 0.84918 | 0.55265 | -0.08687 | -0.51821 |
| Mucus thickness, $M_{\text{thickness}}$ | 0.50251 | 0.45874 | 0.23693 | 0.16992 |

**Figure 8.**
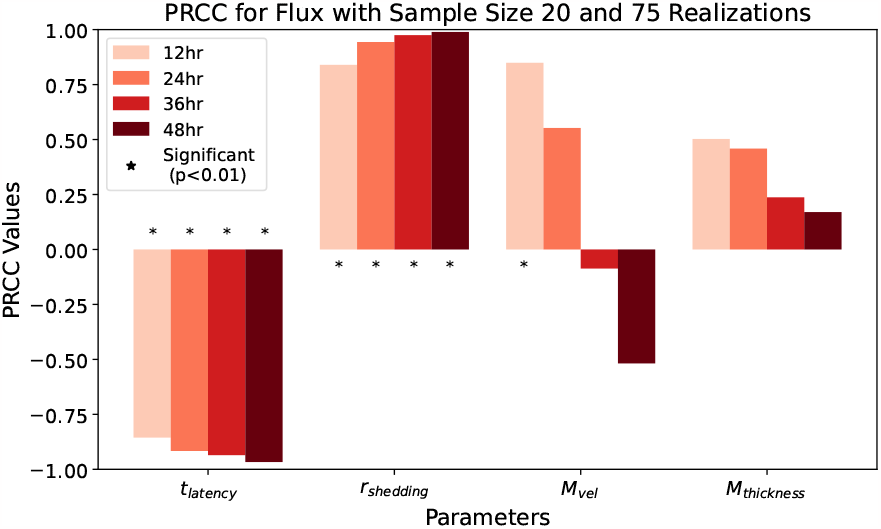
PRCC results for flux at 12, 24, 36, and 48 hours.

Intriguingly, mucus advection velocity starts with a relatively strong positive impact on flux, but we do not detect a significant effect at later time points. We surmise this behavior is a result of the non-monotonicity of the relationship between mucus advection velocity and flux.

Figure 9 and Table 10 show the *virion flux outcomes* at various *M*_vel_ and *M*_thickness_ values while fixing *t*_latency_ = 3 hr and *r*_shedding_ = 100. We see that given any fixed mucus thickness between 12.75 *μ*m and 21.25 *μ*m, the flux outcome values increase and then decrease as advection velocity increases from 36.67 to 220.00 *μ*m/sec. This result confirms that flux is not linearly dependent on *M*_vel_. Hence the PRCC method cannot extract valid information about linear dependency between them.

**Table 10:** Virion flux values (in millions) at 36 hours versus advection velocity and mucus thickness, while fixing *t*_latency_ = 3 hr and *r*_shedding_ = 100 infectious RNA copies per day.

| Flux ($\times 10^6$) | | $M_{\text{thickness}} (\mu\text{m})$ | | | | | |
| --- | --- | --- | --- | --- | --- | --- | --- |
|  |  | 12.75 | 14.45 | 16.15 | 17.85 | 19.55 | 21.25 |
| $M_{\text{vel}} (\mu\text{m/sec})$ | 220.00 | 5.66 | 5.54 | 5.28 | 5.28 | 5.04 | 4.78 |
|  | 183.33 | 6.32 | 6.17 | 5.98 | 6.10 | 5.89 | 5.72 |
|  | 146.67 | 6.70 | 6.52 | 6.72 | 6.55 | 6.52 | 6.45 |
|  | 110.00 | 6.32 | 6.57 | 6.62 | 6.83 | 7.11 | 7.14 |
|  | 73.33 | 4.92 | 5.61 | 6.07 | 6.29 | 6.62 | 6.90 |
|  | 36.67 | 1.42 | 2.02 | 2.74 | 3.29 | 3.81 | 4.31 |

**Figure 9.**
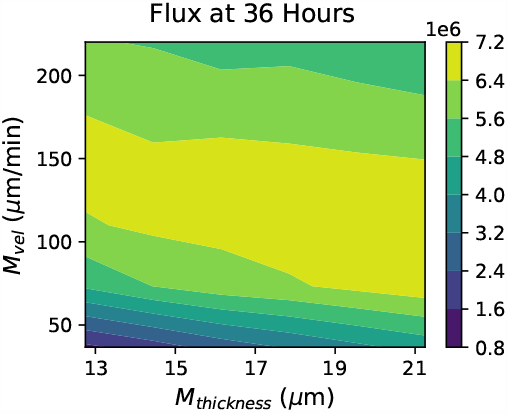
A contour plot showing the 36-hour virion flux outcome for various values of mucus advection velocity and mucus thickness, while fixing *t*_latency_ at 3 hours and *r*_shedding_ = 100 infectious RNA copies per day.

Contour plots showing total viral load, infected cell count, and flux at various *M*_vel_ and *M*_thickness_ values at 12, 24, 36, 48 hours can be found in Appendix C.

## 5 Concluding Remarks

Employing a physiologically faithful, stochastic, spatial model of inhaled SARS-CoV-2 exposure and infection [10], and focusing in the human nasal passage, we expand previous studies [12] in two important directions: (i) incorporation of host variability in physiology (mucus thickness and clearance velocity in the nasal passage), and (ii) application of global sensitivity methods (Partial Rank Correlation Coefficient (PRCC) analysis and Latin Hypercube Sampling (LHS)). In doing so, we decouple potential correlations between physiological parameters and parameters governing virion-cell infection kinetics, thereby isolating outcome sensitivity to each potential driver of heterogeneity. We perform sufficient model simulations to get robust statistics of model-generated outcomes at each sampled location in the parameter space of the model. We then apply PRCC and LHS to evaluate sensitivity and heterogeneity in infection outcomes to within-host variability in nasal mucus thickness and clearance velocity and in virion-cell infection kinetics.

We simulate the progression of infection in the nasal passage within the 48 hours immediately after the infection of one cell at the entrance. We use LHS to select sample points in the four-parameter space: latency time *t*_latency_ from cell infection to extracellular shedding of viral RNA copies, shedding rate of infectious RNA copies *r*_shedding_, mucus advection velocity *M*_vel_, and mucus thickness *M*_thickness_. Then we apply PRCC analysis to analyze the relationship between the parameters and the simulation outcome data: total infectious viral load, infected cell count, and flux of virions out of the nasal passage.

Our results show that throughout the immediate 48 hours post infection, the latency time (*t*_latency_) of newly infected cells has a strong negative PRCC, and the virion shedding rate (*r*_shedding_) has a strong positive PRCC for all three outcome metrics. With total viral load and infected cell count, mucus velocity (*M*_vel_) in the nasal passage has a negative PRCC that becomes more significant over 48 hours. We do not detect a significant effect of mucus thickness (*M*_thickness_) on any of the three outcomes. The unusual progression of PRCC results for *M*_vel_ and viral flux is explained by the non-monotonicity of the relationship between them, which dilutes the conclusions that can be drawn.

The salient insight gained from this study is that the observed population heterogeneity in the early outcomes from nasal exposure and infection to SARS-CoV-2 is reproduced herein, and therefore completely explainable from a mechanistic, physiologically faithful, spatially resolved exposure and infection model. Furthermore, by using global sensitivity analyses, the mechanistic drivers of outcome sensitivity and thereby population heterogeneity can be rank-ordered.

Variability in virus-host cell kinetics, arising from some combination of how the host cells react to exposure and infection by a specific SARS-CoV-2 variant, is the leading-order driver of outcome sensitivity and heterogeneity. Specifically: *Linear variations in the latency time between an infectious virus-cell encounter and onset of extracellular shedding of viral RNA copies and logarithmic variations in the extracellular shedding rate of infectious RNA copies by infected cells are each responsible for exponential heterogeneity in outcomes of nasal viral titers, infected nasal cells, and viral flux from the nasal passage toward the trachea*. We have demonstrated this for latency times between 3 and 9 hours and shedding rates between 10 and 1000 infectious RNA copies / day. Further, these exponential outcome variations arise fast and persist over 48 hours. These results are consistent with the clinical data on nasal swab tests, showing that huge variations in nasal titers from positive tests are feasible even for persons infected for as few as 12 or 24 hours.

*Variation in mucus clearance velocity has a time-dependent influence on outcomes: very weak in the first 24 hours post infection, then growing to a significant impact on outcomes in the subsequent 24 hours. This result suggests that the slowdown in mucus velocity post infection likely takes 12-24 hours to take effect, but once mucus advection slows in day 2, it amplifies viral load and infected cell count. The thickness of the nasal mucus layer has a minor impact on infection outcomes*.

These results and insights strongly suggest the need for experimental data to be collected spanning different variants of SARS-CoV-2, spanning nasal cultures grown from a diverse collection of individuals, and then careful measurements of the mechanistic parameters in our model. We note that cell culture experiments must be focused on measurements of infection probability per virus-cell encounter, latency time, and extracellular shedding rate. The outcome metrics of total shed viral load and number of infected cells in a cell culture will not be representative of in vivo nasal infection, since there is no mucus clearance in cell cultures. In order for these insights to be “actionable” for medical treatment, an immediate nasal culture can determine the virus-cell infection kinetics of an individual, and single-cell measurements of latency time and replication rate would guide the need for rapid drug or antiviral therapies applied directly to the nasal passage. Lastly, the flexibility and robustness of our model and simulation platform are adaptable for future investigations into other respiratory viruses.

## Author Contributions

Conceptualization, M.G.F., A.C., A.A., T.W., L.Z. and H.C.; methodology, M.G.F., T.W., L.Z. and H.C.; software, A.C., J.P., K.M., L.Z. and H.C.; validation and formal analysis, L.Z., H.C. and M.G.F.; writing—original draft preparation, L.Z. and M.G.F.; writing—review and editing, all authors; project administration, M.G.F. All authors have read and agreed to the submitted version of the manuscript.

## Data Availability

All data is available upon request to the senior author, M.G.F.

## Appendix A

**Figure 10.**
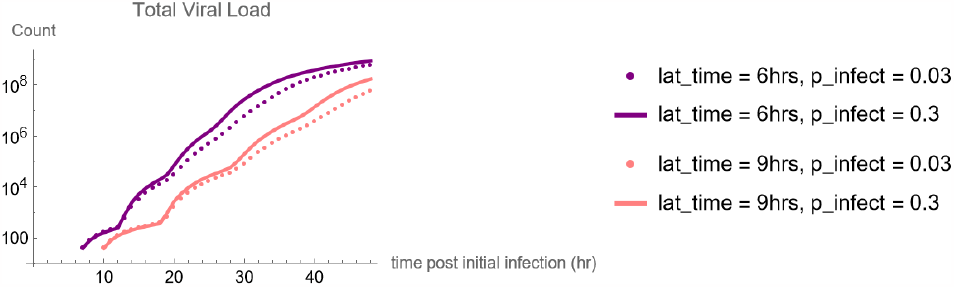
The total viral load in the nasal passage during the 48 hours post infection, given different latency times and the probabilities of infection per virion-cell encounter per second.

For latency time at 6 and 9 hours, we use Eq. 9 compute the percent difference between the total loads during the 48 hours post infection given probabilities of infection per virion-cell encounter per second (*p*_infect_) at 0.03 versus 0.3.

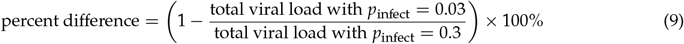

In each case, whether the latency time is 6 or 9 hours, we change the probability to infect by one order of magnitude, but the total viral loads after 48 hours only differ by a multiplicative factor.

**Figure 11.**
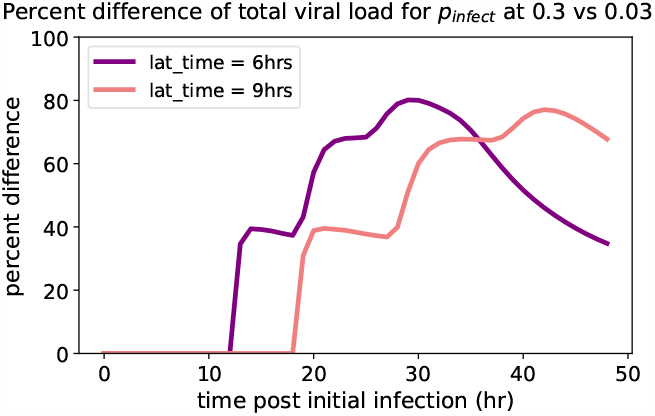
The percent difference between the total viral loads during the 48 hours post infection given different probabilities of infection per virion-cell encounter per second.

## Appendix B

**Figure 12.**
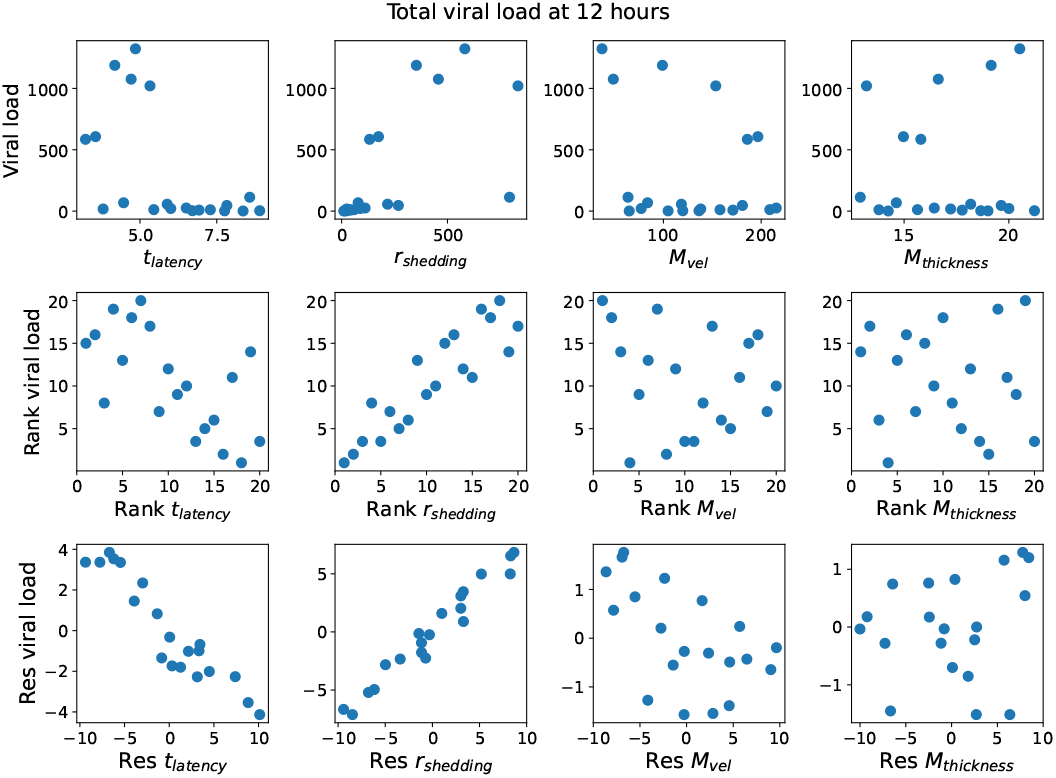
Each row from top to bottom: (1) total viral load at 12 hours versus each parameter; (2) ranks of total viral load at 12 hours versus ranks of each parameter; (3) residuals of total viral load at 12 hours versus the residuals of each parameter.

**Figure 13.**
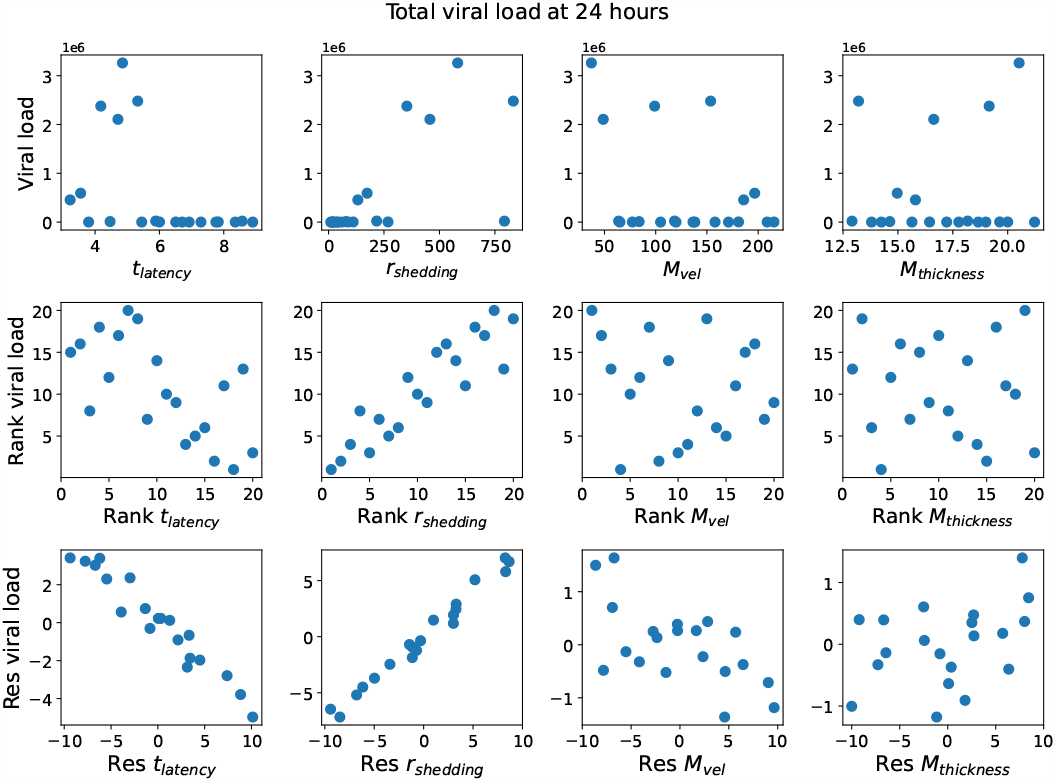
Each row from top to bottom: (1) total viral load at 24 hours versus each parameter; (2) ranks of total viral load at 24 hours versus ranks of each parameter; (3) residuals of total viral load at 24 hours versus the residuals of each parameter.

**Figure 14.**
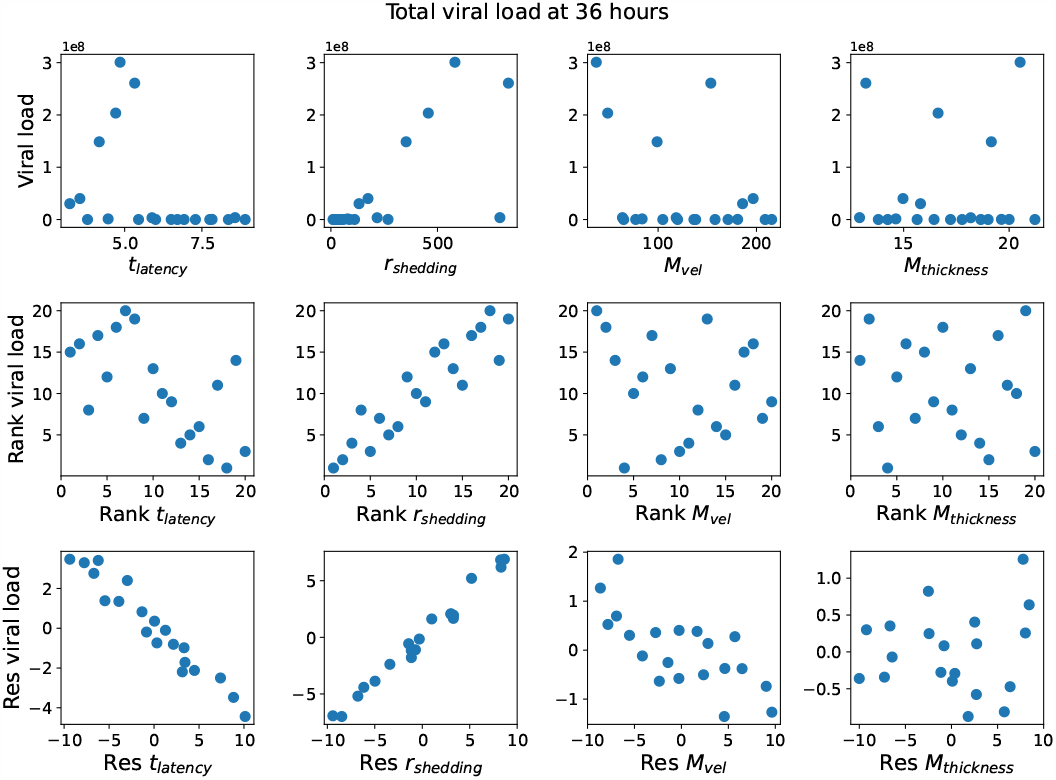
Each row from top to bottom: (1) total viral load at 36 hours versus each parameter; (2) ranks of total viral load at 36 hours versus ranks of each parameter; (3) residuals of total viral load at 36 hours versus the residuals of each parameter.

**Figure 15.**
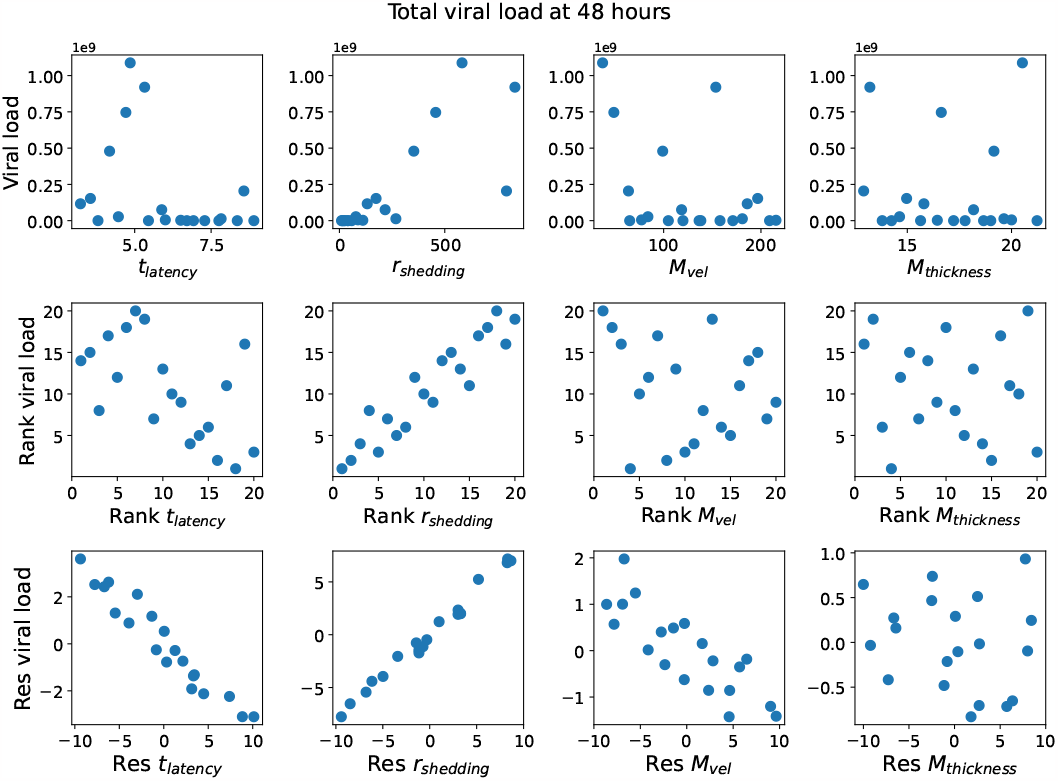
Each row from top to bottom: (1) total viral load at 48 hours versus each parameter; (2) ranks of total viral load at 48 hours versus ranks of each parameter; (3) residuals of total viral load at 48 hours versus the residuals of each parameter.

**Figure 16.**
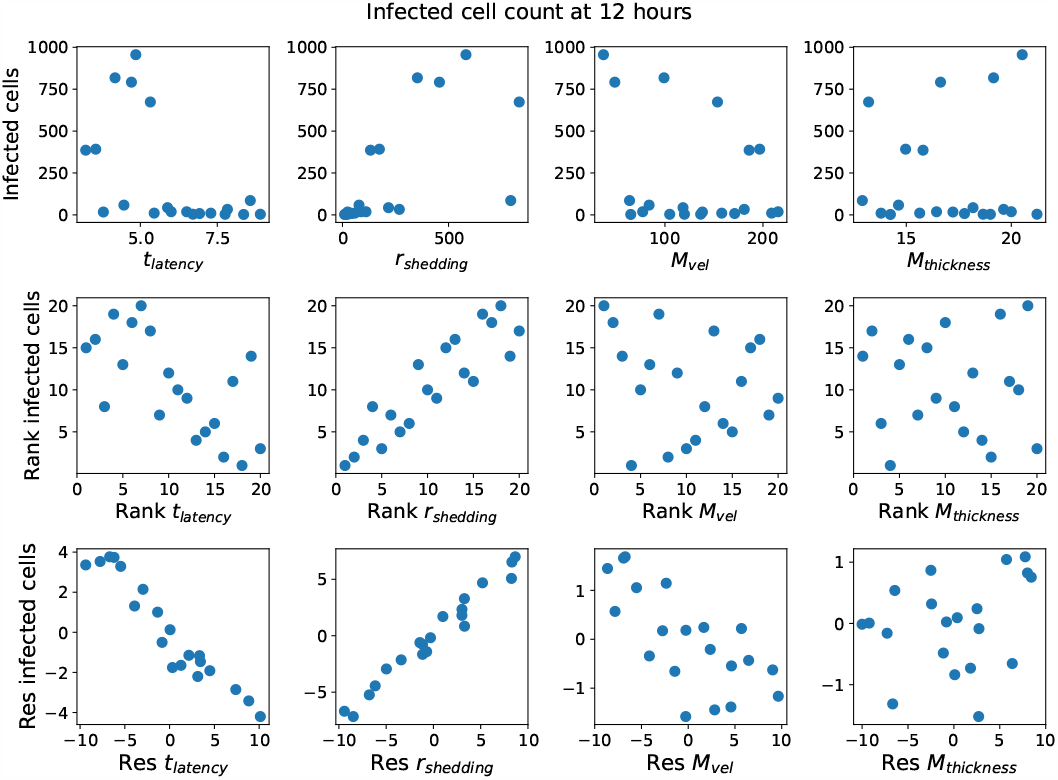
Each row from top to bottom: (1) infected cell count at 12 hours versus each parameter; (2) ranks of infected cell count at 12 hours versus ranks of each parameter; (3) residuals of infected cell count at 12 hours versus the residuals of each parameter.

**Figure 17.**
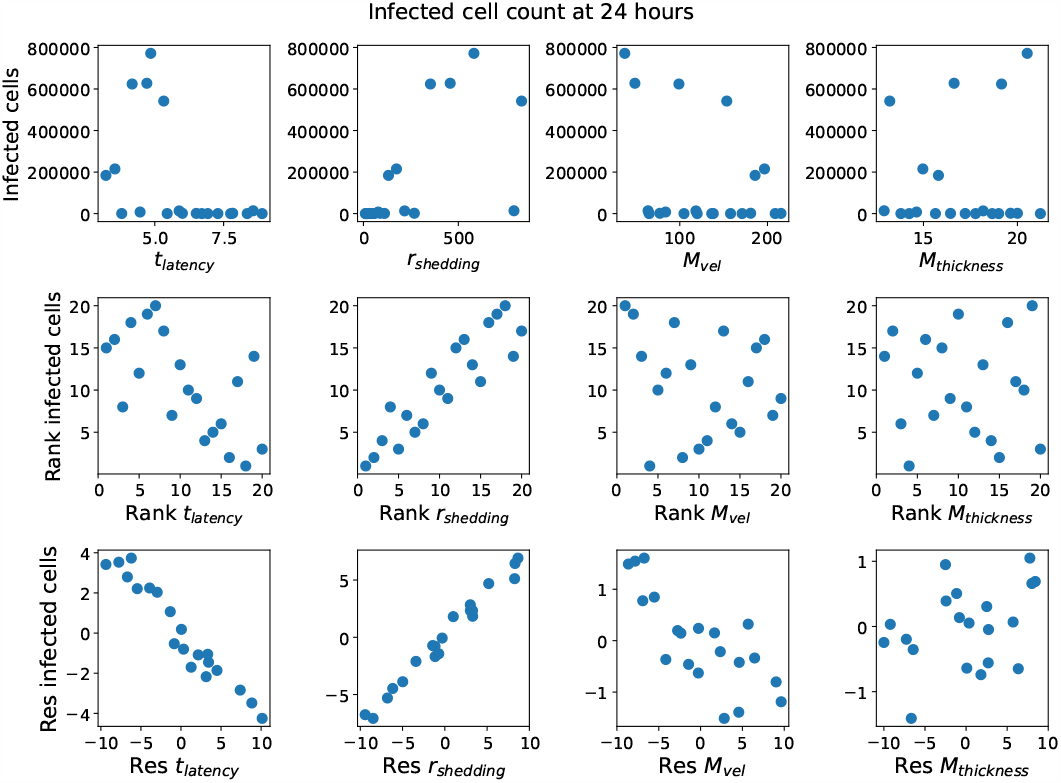
Each row from top to bottom: (1) infected cell count at 24 hours versus each parameter; (2) ranks of infected cell count at 24 hours versus ranks of each parameter; (3) residuals of infected cell count at 24 hours versus the residuals of each parameter.

**Figure 18.**
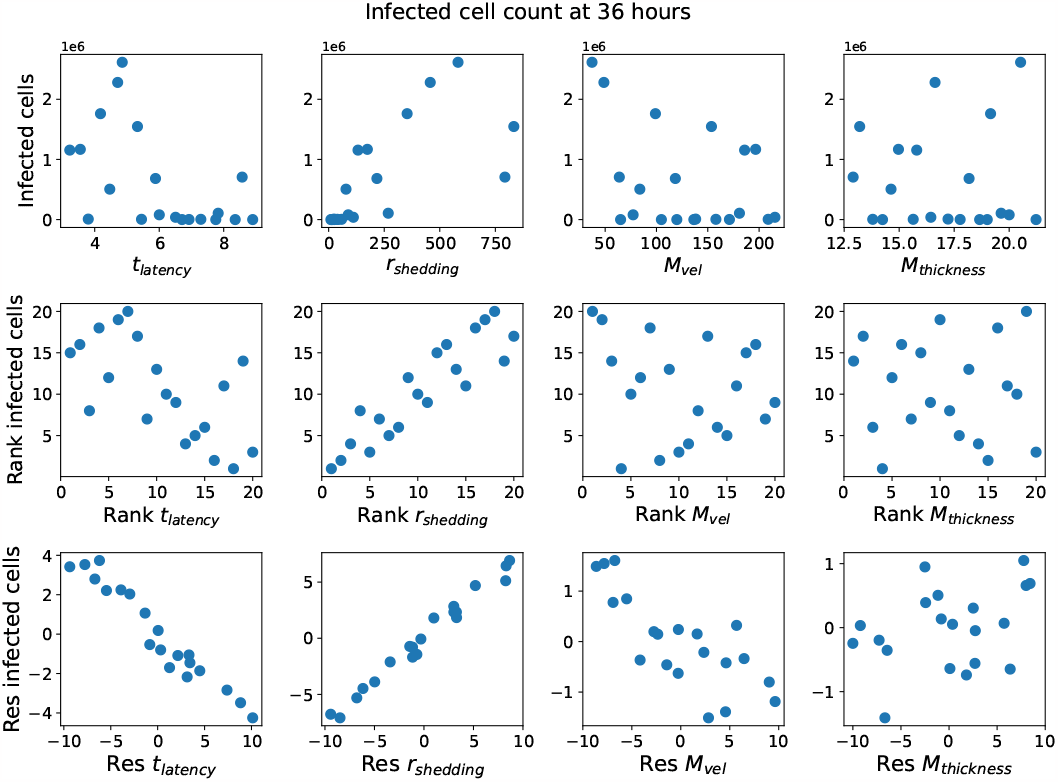
Each row from top to bottom: (1) infected cell count at 36 hours versus each parameter; (2) ranks of infected cell count at 36 hours versus ranks of each parameter; (3) residuals of infected cell count at 36 hours versus the residuals of each parameter.

**Figure 19.**
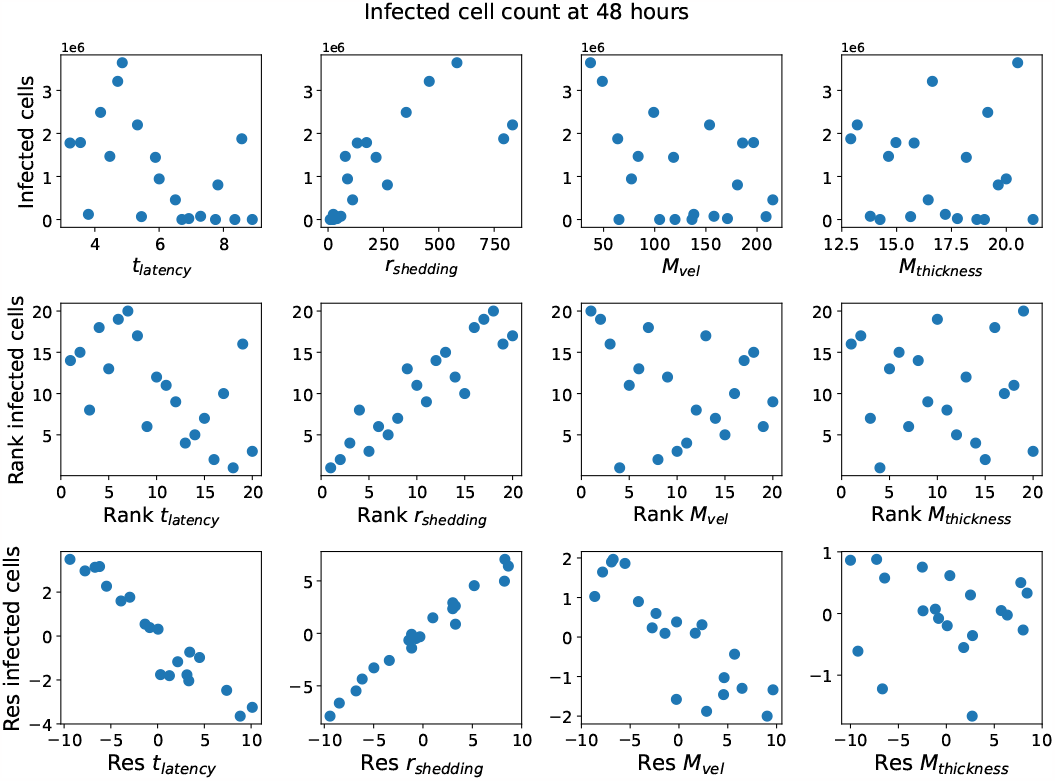
Each row from top to bottom: (1) infected cell count at 48 hours versus each parameter; (2) ranks of infected cell count at 48 hours versus ranks of each parameter; (3) residuals of infected cell count at 48 hours versus the residuals of each parameter.

**Figure 20.**
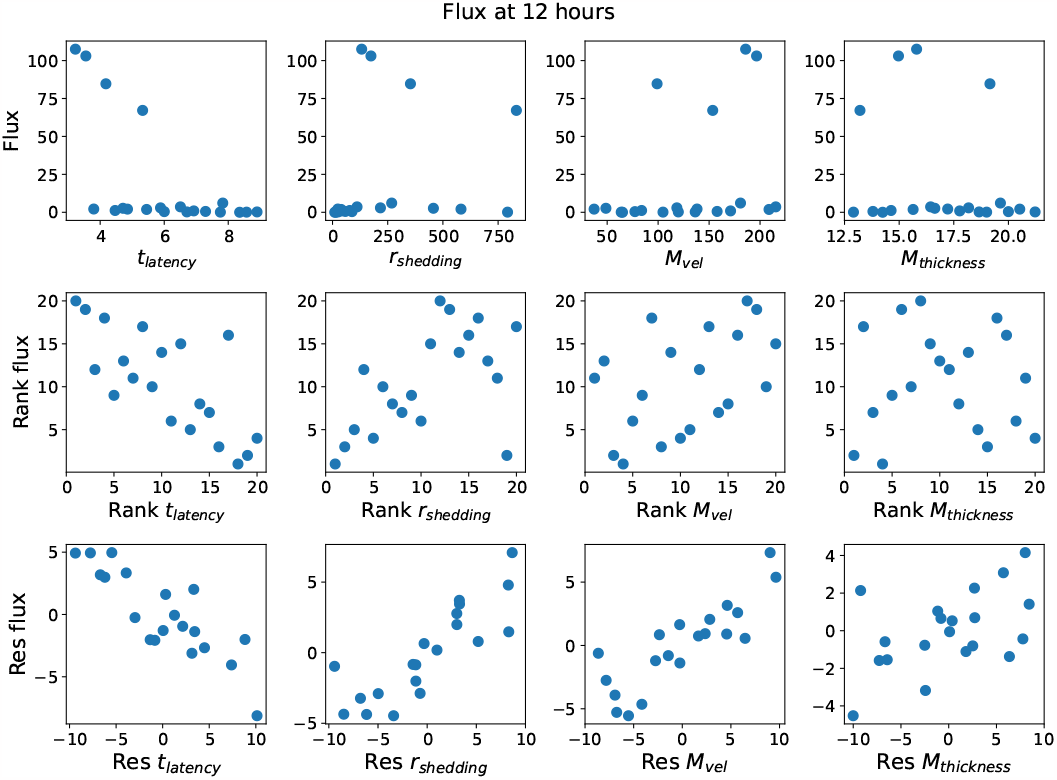
Each row from top to bottom: (1) flux at 12 hours versus each parameter; (2) ranks of flux at 12 hours versus ranks of each parameter; (3) residuals of flux at 12 hours versus the residuals of each parameter.

**Figure 21.**
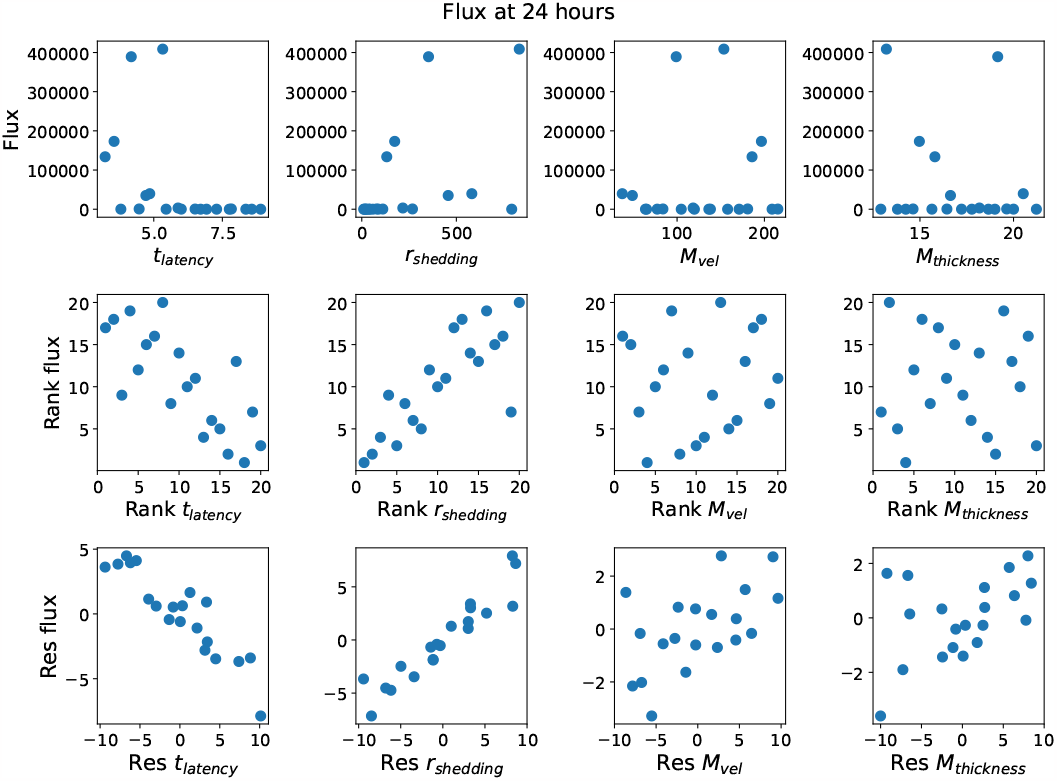
Each row from top to bottom: (1) flux at 24 hours versus each parameter; (2) ranks of flux at 24 hours versus ranks of each parameter; (3) residuals of flux at 24 hours versus the residuals of each parameter.

**Figure 22.**
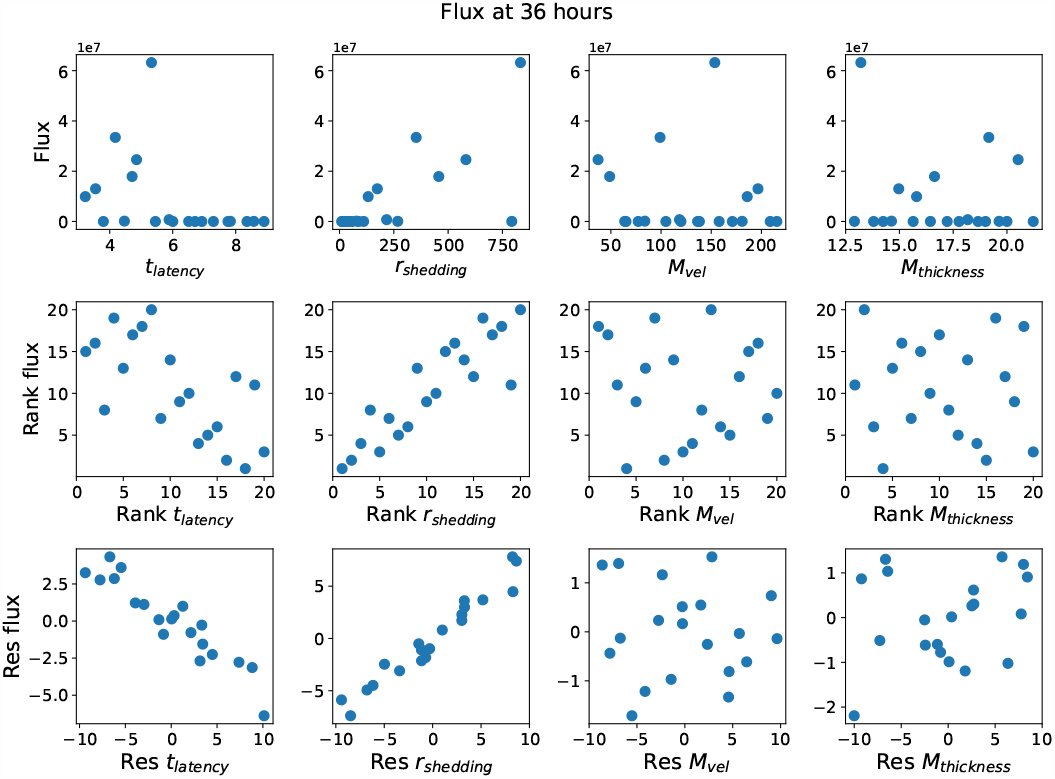
Each row from top to bottom: (1) flux at 36 hours versus each parameter; (2) ranks of flux at 36 hours versus ranks of each parameter; (3) residuals of flux at 36 hours versus the residuals of each parameter.

**Figure 23.**
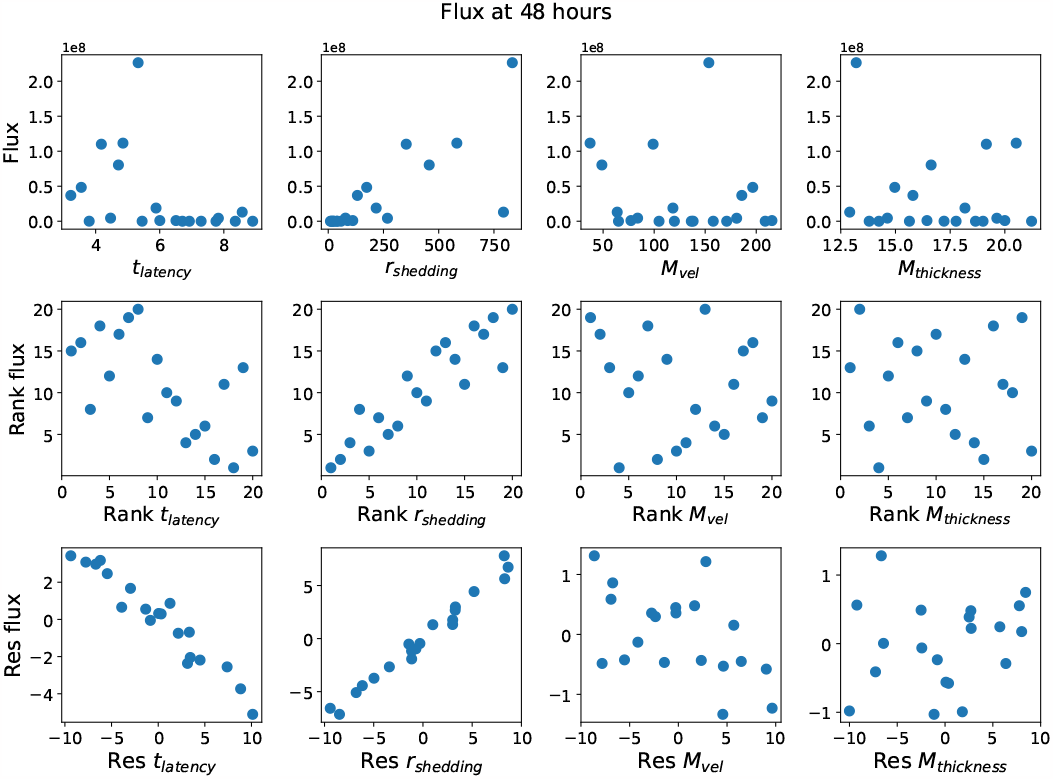
Each row from top to bottom: (1) flux at 48 hours versus each parameter; (2) ranks of flux at 48 hours versus ranks of each parameter; (3) residuals of flux at 48 hours versus the residuals of each parameter.

## APPENDIX C

**Figure 24.**
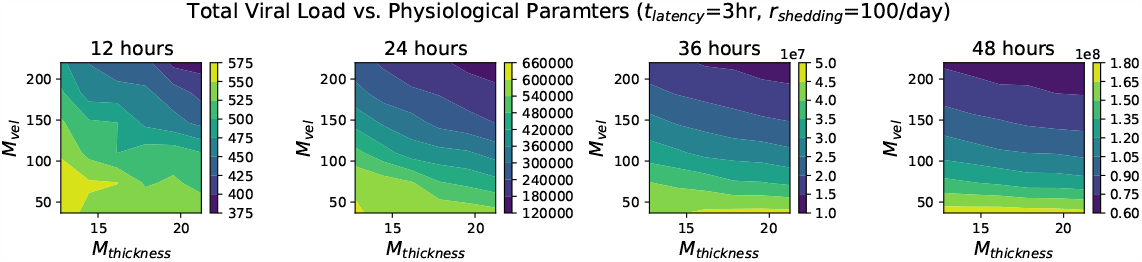
A contour plot showing the total viral load at 12, 24, 36, and 48 hours for various values of mucus advection velocity and mucus thickness, while fixing *t*_latency_ = 3 hours and *r*_shedding_ = 100 infectious RNA copies per day.

**Figure 25.**
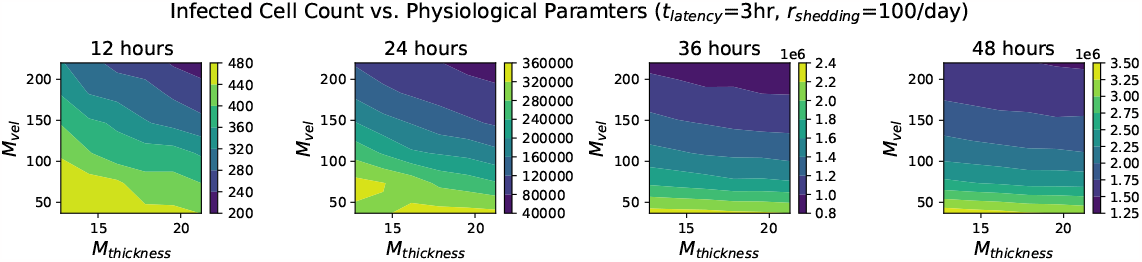
A contour plot showing the infected cell count at 12, 24, 36, and 48 hours for various values of mucus advection velocity and mucus thickness, while fixing *t*_latency_ = 3 hours and *r*_shedding_ = 100 infectious RNA copies per day.

**Figure 26.**
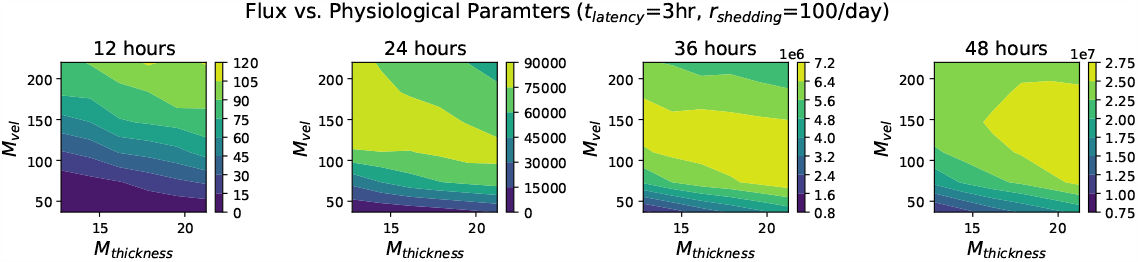
A contour plot showing the flux at 12, 24, 36, and 48 hours for various values of mucus advection velocity and mucus thickness, while fixing *t*_latency_ = 3 hours and *r*_shedding_ = 100 infectious RNA copies per day.

During the first 12 hours, virions can easily reach uninfected cells via (1) diffusion in the PCL only, or (2) a short trip in the mucus layer and diffusion in PCL. Increasing the advection velocity and mucus thickness do not affect (1). Faster advection will flush out more virions that enter the mucus layer, but virions do not have to stay in the mucus layer for long to infect as in (2), and faster advection won’t affect these virions as much.

At later times, however, most cells that can be reached via (1) and (2) have already been infected, so virions have to travel further to find uninfected cells, especially in the azimuthal direction perpendicular to advection, which can only be achieved via diffusion. That requires the virions to spend longer in the ASL layers but that increases the probability of them getting deeper into the mucus layer. In this case, faster advection increases the probability of those virions being carried out of the generation before they have a chance to re-enter the PCL.

The impact of mucus layer thickness might depend more on the values of other parameters, e.g. advection velocity. Consider the following scenarios: (i) Early on, virions are able to infect new cells without being in the mucus layer for a long time. In this case, increasing the mucus layer thickness likely doesn’t interfere with the spread of infection very much. (ii) At later times, if advection velocity is high, then virions are likely to be flushed out before they have time to diffuse deeply enough into the mucus layer for the mucus layer thickness to matter. (iii) At later times, if advection velocity is low, then virions can stay in the mucus layer for a longer period of time, which allows them to diffuse further in the azimuthal direction and for them to re-enter the PCL layer to infect cells. Meanwhile a thicker mucus layer gives virions the room to diffuse further away from the epithelium cells which further increases the time virions can spend in the mucus layer. In combination, the low advection velocity and higher mucus layer thickness may allow virions to travel further and spread infection.

## References

[1] Roman Wölfel, Victor M Corman, Wolfgang Guggemos, Michael Seilmaier, Sabine Zange, Marcel A Müller, Daniela Niemeyer, Terry C Jones, Patrick Vollmar, Camilla Rothe, et al. Virological assessment of hospitalized patients with covid-2019. Nature, 581(7809):465–469, 2020.

[2] Ashish Goyal, E. Fabian Cardozo-Ojeda, and Joshua T. Schiffer. Potency and timing of antiviral therapy as determinants of duration of sars-cov-2 shedding and intensity of inflammatory response. Science Advances, 6(47):eabc7112, 2020.

[3] Nadège Néant, Guillaume Lingas, Quentin Le Hingrat, Jade Ghosn, Ilka Engelmann, Quentin Lepiller, Alexandre Gaymard, Virginie Ferré, Cédric Hartard, Jean-Christophe Plantier, Vincent Thibault, Julien Marlet, Brigitte Montes, Kevin Bouiller, François-Xavier Lescure, Jean-François Timsit, Emmanuel Faure, Julien Poissy, Christian Chidiac, François Raffi, Antoine Kimmoun, Manuel Etienne, Jean-Christophe Richard, Pierre Tattevin, Denis Garot, Vincent Le Moing, Delphine Bachelet, Coralie Tardivon, Xavier Duval, Yazdan Yazdanpanah, France Mentré, Cédric Laouénan, Benoit Visseaux, Jérémie Guedj, for the French COVID Cohort Investigators, and French Cohort Study groups. Modeling sars-cov-2 viral kinetics and association with mortality in hospitalized patients from the french covid cohort. Proceedings of the National Academy of Sciences, 118(8):e2017962118, 2021.

[4] Elisa Teyssou, Héloise Delagrèverie, Benoit Visseaux, Sidonie Lambert-Niclot, Ségolène Brichler, Valentine Ferre, Stéphane Marot, Aude Jary, Eve Todesco, Aurélie Schnuriger, Emna Ghidaoui, Basma Abdi, Sepideh Akhavan, Nadhira Houhou-Fidouh, Charlotte Charpentier, Laurence Morand-Joubert, David Boutolleau, Diane Descamps, Vincent Calvez, Anne Geneviève Marcelin, and Cathia Soulie. The delta sars-cov-2 variant has a higher viral load than the beta and the historical variants in nasopharyngeal samples from newly diagnosed covid-19 patients. Journal of Infection, 83(4):e1–e3, 2021.

[5] Ruian Ke, Carolin Zitzmann, David D. Ho, Ruy M. Ribeiro, and Alan S. Perelson. In vivo kinetics of sars-cov-2 infection and its relationship with a person’s infectiousness. Proceedings of the National Academy of Sciences, 118(49):e2111477118, 2021.

[6] Baisheng Li, Aiping Deng, Kuibiao Li, Yao Hu, Zhencui Li, Yaling Shi, Qianling Xiong, Zhe Liu, Qianfang Guo, Lirong Zou, et al. Viral infection and transmission in a large, well-traced outbreak caused by the sars-cov-2 delta variant. Nature communications, 13(1):460, 2022.

[7] Alexandre Bolze, Shishi Luo, Simon White, Elizabeth T. Cirulli, Dana Wyman, Andrew Dei Rossi, Henrique Machado, Tyler Cassens, Sharoni Jacobs, Kelly M. Schiabor Barrett, Francisco Tanudjaja, Kevin Tsan, Jason Nguyen, Jimmy M. Ramirez, Efren Sandoval, Xueqing Wang, David Wong, David Becker, Marc Laurent, James T. Lu, Magnus Isaksson, Nicole L. Washington, and William Lee. Sars-cov-2 variant delta rapidly displaced variant alpha in the united states and led to higher viral loads. Cell Reports Medicine, 3(3):100564, 2022.

[8] Ruian Ke, Pamela P Martinez, Rebecca L Smith, Laura L Gibson, Agha Mirza, Madison Conte, Nicholas Gallagher, Chun Huai Luo, Junko Jarrett, Ruifeng Zhou, et al. Daily longitudinal sampling of sars-cov-2 infection reveals substantial heterogeneity in infectiousness. Nature microbiology, 7(5):640–652, 2022.

[9] Miguel Garcia-Knight, Khamal Anglin, Michel Tassetto, Scott Lu, Amethyst Zhang, Sarah A. Goldberg, Adam Catching, Michelle C. Davidson, Joshua R. Shak, Mariela Romero, Jesus Pineda-Ramirez, Ruth Diaz-Sanchez, Paulina Rugart, Kevin Donohue, Jonathan Massachi, Hannah M. Sans, Manuella Djomaleu, Sujata Mathur, Venice Servellita, David McIlwain, Brice Gaudiliere, Jessica Chen, Enrique O. Martinez, Jacqueline M. Tavs, Grace Bronstone, Jacob Weiss, John T. Watson, Melissa Briggs-Hagen, Glen R. Abedi, George W. Rutherford, Steven G. Deeks, Charles Chiu, Sharon Saydah, Michael J. Peluso, Claire M. Midgley, Jeffrey N. Martin, Raul Andino, and J. Daniel Kelly. Infectious viral shedding of sars-cov-2 delta following vaccination: A longitudinal cohort study. PLOS Pathogens, 18(9):1–17, 09 2022.

[10] Alexander Chen, Timothy Wessler, Katherine Daftari, Kameryn Hinton, Richard C. Boucher, Raymond Pickles, Ronit Freeman, Samuel K. Lai, and M. Gregory Forest. Modeling insights into sars-cov-2 respiratory tract infections prior to immune protection. Biophysical Journal, 121(9):1619–1631, 2022.

[11] Andreas C. Aristotelous, Alex Chen, and M. Gregory Forest. A hybrid discrete-continuum model of immune responses to sars-cov-2 infection in the lung alveolar region, with a focus on interferon induced innate response. Journal of Theoretical Biology, 555:111293, 2022.

[12] Jason Pearson, Timothy Wessler, Alex Chen, Richard C. Boucher, Ronit Freeman, Samuel K. Lai, Raymond Pickles, and M. Gregory Forest. Modeling identifies variability in sars-cov-2 uptake and eclipse phase by infected cells as principal drivers of extreme variability in nasal viral load in the 48 h post infection. Journal of Theoretical Biology, 565:111470, 2023.

[13] Feng Guo, Sixing Li, Mehmet Umut Caglar, Zhangming Mao, Wu Liu, Andrew Woodman, Jamie J. Arnold, Claus O. Wilke, Tony Jun Huang, and Craig E. Cameron. Single-cell virology: On-chip investigation of viral infection dynamics. Cell Reports, 21(6):1692–1704, 2017.

[14] Wu Liu, Mehmet U. Caglar, Zhangming Mao, Andrew Woodman, Jamie J. Arnold, Claus O. Wilke, and Craig E. Cameron. More than efficacy revealed by single-cell analysis of antiviral therapeutics. Science Advances, 5(10):eaax4761, 2019.

[15] Jeffrey Y Lee, Peter AC Wing, Dalia S Gala, Marko Noerenberg, Aino I Järvelin, Joshua Titlow, Xiaodong Zhuang, Natasha Palmalux, Louisa Iselin, Mary Kay Thompson, Richard M Parton, Maria Prange-Barczynska, Alan Wainman, Francisco J Salguero, Tammie Bishop, Daniel Agranoff, William James, Alfredo Castello, Jane A McKeating, and Ilan Davis. Absolute quantitation of individual sars-cov-2 rna molecules provides a new paradigm for infection dynamics and variant differences. eLife, 11:e74153. jan 2022.

[16] Qian Li, Kadambari Vijaykumar, Scott E. Phillips, Shah S. Hussain, Nha V. Huynh, Courtney M. Fernandez-Petty, Jacelyn E. Peabody Lever, Jeremy B. Foote, Janna Ren, Javier Campos-Gómez, Farah Abou Daya, Nathaniel W. Hubbs, Harrison Kim, Ezinwanne Onuoha, Evan R. Boitet, Lianwu Fu, Hui Min Leung, Linhui Yu, Thomas W. Detchemendy, Levi T. Schaefers, Jennifer L. Tipper, Lloyd J. Edwards, Sixto M. Leal Jr., Kevin S. Harrod, Guillermo J. Tearney, and Steven M. Rowe. Mucociliary transport deficiency and disease progression in syrian hamsters with sars-cov-2 infection. JCI Insight, 8(1), 1 2023.

[17] Chris Davis, Nicola Logan, Grace Tyson, Richard Orton, William T. Harvey, Jonathan S. Perkins, Guy Mollett, Rachel M. Blacow, The COVID-19 Genomics UK (COG-UK) Consortium, Thomas P. Peacock, Wendy S. Barclay, Peter Cherepanov, Massimo Palmarini, Pablo R. Murcia, Arvind H. Patel, David L. Robertson, John Haughney, Emma C. Thomson, Brian J. Willett, and on behalf of the COVID-19 DeplOyed VaccinE (DOVE) Cohort Study investigators. Reduced neutralisation of the delta (b.1.617.2) sars-cov-2 variant of concern following vaccination. PLOS Pathogens, 17(12):1–15, 12 2021.

[18] Emma C Wall, Mary Wu, Ruth Harvey, Gavin Kelly, Scott Warchal, Chelsea Sawyer, Rodney Daniels, Philip Hobson, Emine Hatipoglu, Yenting Ngai, Saira Hussain, Jerome Nicod, Robert Goldstone, Karen Ambrose, Steve Hindmarsh, Rupert Beale, Andrew Riddell, Steve Gamblin, Michael Howell, George Kassiotis, Vincenzo Libri, Bryan Williams, Charles Swanton, Sonia Gandhi, and David LV Bauer. Neutralising antibody activity against sars-cov-2 vocs b.1.617.2 and b.1.351 by bnt162b2 vaccination. The Lancet, 397(10292):2331–2333, 2021.

[19] Emma C Wall, Mary Wu, Ruth Harvey, Gavin Kelly, Scott Warchal, Chelsea Sawyer, Rodney Daniels, Lorin Adams, Philip Hobson, Emine Hatipoglu, Yenting Ngai, Saira Hussain, Karen Ambrose, Steve Hindmarsh, Rupert Beale, Andrew Riddell, Steve Gamblin, Michael Howell, George Kassiotis, Vincenzo Libri, Bryan Williams, Charles Swanton, Sonia Gandhi, and David LV Bauer. Azd1222-induced neutralising antibody activity against sars-cov-2 delta voc. The Lancet, 398(10296):207–209, 2021.

[20] Katrina A. Lythgoe, Matthew Hall, Luca Ferretti, Mariateresa de Cesare, George MacIntyre-Cockett, Amy Trebes, Monique Andersson, Newton Otecko, Emma L. Wise, Nathan Moore, Jessica Lynch, Stephen Kidd, Nicholas Cortes, Matilde Mori, Rebecca Williams, Gabrielle Vernet, Anita Justice, Angie Green, Samuel M. Nicholls, M. Azim Ansari, Lucie Abeler-Dörner, Catrin E. Moore, Timothy E. A. Peto, David W. Eyre, Robert Shaw, Peter Simmonds, David Buck, John A. Todd, on behalf of the Oxford Virus Sequencing Analysis Group (OVSG)‡, Thomas R. Connor, Shirin Ashraf, Ana da Silva Filipe, James Shepherd, Emma C. Thomson, The COVID-19 Genomics UK (COG-UK) Consortium§, David Bonsall, Christophe Fraser, and Tanya Golubchik. Sars-cov-2 within-host diversity and transmission. Science, 372(6539):eabg0821, 2021.

[21] Jamie Lopez Bernal, Nick Andrews, Charlotte Gower, Eileen Gallagher, Ruth Simmons, Simon Thelwall, Julia Stowe, Elise Tessier, Natalie Groves, Gavin Dabrera, Richard Myers, Colin N.J. Campbell, Gayatri Amirthalingam, Matt Edmunds, Maria Zambon, Kevin E. Brown, Susan Hopkins, Meera Chand, and Mary Ramsay. Effectiveness of covid-19 vaccines against the b.1.617.2 (delta) variant. New England Journal of Medicine, 385(7):585–594, 2021. PMID: 34289274.

[22] Tyler N. Starr, Allison J. Greaney, Amin Addetia, William W. Hannon, Manish C. Choudhary, Adam S. Dingens, Jonathan Z. Li, and Jesse D. Bloom. Prospective mapping of viral mutations that escape antibodies used to treat covid-19. Science, 371(6531):850–854, 2021.

[23] Emma C. Thomson, Laura E. Rosen, James G. Shepherd, Roberto Spreafico, Ana da Silva Filipe, Jason A. Wojcechowskyj, Chris Davis, Luca Piccoli, David J. Pascall, Josh Dillen, Spyros Lytras, Nadine Czudnochowski, Rajiv Shah, Marcel Meury, Natasha Jesudason, Anna De Marco, Kathy Li, Jessica Bassi, Aine O’Toole, Dora Pinto, Rachel M. Colquhoun, Katja Culap, Ben Jackson, Fabrizia Zatta, Andrew Rambaut, Stefano Jaconi, Vattipally B. Sreenu, Jay Nix, Ivy Zhang, Ruth F. Jarrett, William G. Glass, Martina Beltramello, Kyriaki Nomikou, Matteo Pizzuto, Lily Tong, Elisabetta Cameroni, Tristan I. Croll, Natasha Johnson, Julia Di Iulio, Arthur Wickenhagen, Alessandro Ceschi, Aoife M. Harbison, Daniel Mair, Paolo Ferrari, Katherine Smollett, Federica Sallusto, Stephen Carmichael, Christian Garzoni, Jenna Nichols, Massimo Galli, Joseph Hughes, Agostino Riva, Antonia Ho, Marco Schiuma, Malcolm G. Semple, Peter J.M. Openshaw, Elisa Fadda, J. Kenneth Baillie, John D. Chodera, Suzannah J. Rihn, Samantha J. Lycett, Herbert W. Virgin, Amalio Telenti, Davide Corti, David L. Robertson, and Gyorgy Snell. Circulating sars-cov-2 spike n439k variants maintain fitness while evading antibody-mediated immunity. Cell, 184(5):1171–1187.e20, 2021.

[24] Brian J. Willett, Joe Grove, Oscar A. MacLean, Craig Wilkie, Giuditta De Lorenzo, Wilhelm Furnon, Diego Cantoni, Sam Scott, Nicola Logan, Shirin Ashraf, and et al. Sars-cov-2 omicron is an immune escape variant with an altered cell entry pathway. Nature Microbiology, 7(8):1161–1179, 2022.

[25] Julie Boucau, Caitlin Marino, James Regan, Rockib Uddin, Manish C. Choudhary, James P. Flynn, Geoffrey Chen, Ashley M. Stuckwisch, Josh Mathews, May Y. Liew, Arshdeep Singh, Taryn Lipiner, Autumn Kittilson, Meghan Melberg, Yijia Li, Rebecca F. Gilbert, Zahra Reynolds, Surabhi L. Iyer, Grace C. Chamberlin, Tammy D. Vyas, Marcia B. Goldberg, Jatin M. Vyas, Jonathan Z. Li, Jacob E. Lemieux, Mark J. Siedner, and Amy K. Barczak. Duration of shedding of culturable virus in sars-cov-2 omicron (ba.1) infection. New England Journal of Medicine, 387(3):275–277, 2022. PMID: 35767428.

[26] Michael R. Knowles and Richard C. Boucher. Mucus clearance as a primary innate defense mechanism for mammalian airways. The Journal of Clinical Investigation, 109(5):571–577, 3 2002.

[27] Simeone Marino, Ian B. Hogue, Christian J. Ray, and Denise E. Kirschner. A methodology for performing global uncertainty and sensitivity analysis in systems biology. Journal of Theoretical Biology, 254(1):178–196, 2008.

